# Differential immune gene expression associated with contemporary range expansion of two invasive rodents in Senegal

**DOI:** 10.1101/442160

**Authors:** Nathalie Charbonnel, Maxime Galan, Caroline Tatard, Anne Loiseau, Christophe Amidi Diagne, Ambroise Dalecky, Hugues Parrinello, Stéphanie Rialle, Dany Severac, Carine Brouat

## Abstract

Biological invasions are major anthropogenic changes associated with threats to biodiversity and health. What determines the successful establishment and spread of introduced populations still remains unsolved. Here, we explore the assertion that invasion success relies on immune phenotypic traits that would be advantageous in recently invaded sites. We compared gene expression profiles between anciently and recently established populations of two major invading species, the house mouse *Mus musculus domesticus* and the black rat *Rattus rattus*, in Senegal (West Africa). Transcriptome analyses revealed respectively 364 and 83 differentially expressed genes between anciently and recently established mouse and rat populations. Among them, 20.0% and 10.6% were annotated with functions related to immunity. All immune-related genes detected along the mouse invasion route were over-expressed in recently invaded sites. Genes of the complement activation pathway were over-represented. Results were less straightforward for the black rat as no particular immunological process was over-represented. We revealed changes in transcriptome profiles along invasion routes, although specific patterns differed between the two invasive species. These changes could potentially be driven by increased infection risks in recently invaded sites for the house mouse and stochastic events associated with colonization history for the black rat. These results provide a first step in identifying the immune eco-evolutionary processes potentially involved in invasion success.

## Introduction

Biological invasions are considered as a major component of global change growing threats to biodiversity ^1^, ecosystem functioning^2^ and human health^3^. They may also lead to huge negative economic impacts ^4^. There is hence an urgent need for understanding what determines invasion success (the successful introduction, establishment and spread of species outside their native range) to design effective prevention, control and management strategies. From an eco-evolutionary perspective, invasion success may rely on pre-adaptation within the original range ^5,6^ or on the rapid changes in phenotypic traits (either through selection on new/ standing genetic variation or through adaptive plasticity) that would be advantageous in newly colonized areas ^5,7^. In this context, a corpus of hypotheses has focused on traits that are related with the ability to cope with novel environments ^8^. In particular, important changes in parasite pressures ^8,9^ can occur on introduced species along invasion gradients, from native to invaded areas and within invaded areas from introduction areas to invasion fronts (enemy release, spill-over of native parasites ^10-12^). Thus, invasion success could rely on immune strategies. Two general hypotheses have been developed according to the life history theory. The “evolution of increased competitive ability” (EICA) ^13^ is based on trade-offs between immune traits and other life history traits. It proposes that the loss of parasites that often occurs during the course of invasion should release energetic resources dedicated to defences against parasites. The reallocation of this energy towards other life history traits favouring range expansion (dispersal, reproduction…) should hence be favoured. The “EICA-refined” hypothesis ^14^ considers the possibility that invaders encounter new infection risks during the course of invasion, e.g., through spill-over of native parasites. This hypothesis is based on trade-offs between immune pathways, as these latter are supposed to have different developmental, maintenance and use costs ^15^. This hypothesis therefore suggests that energy allocation to defence against parasites should be modulated in favour of defence (including immune) strategies with lower costs. Considering vertebrates, costly immune pathways such as inflammation could be dampened while less costly ones - for example antibody-mediated responses-could be favoured^14,16^. A recent analysis reviewing empirical studies has shown that in most cases, variations in immune responses were observed during invasions although no general pattern could be emphasized to corroborate the EICA hypotheses ^16^. One of the problems pointed out by the authors was the lack of studies providing a large array of immune responses to properly assess immunocompetence and estimate immune trade-offs. Unfortunately, immune phenotyping often relies on a small number of immune effectors, because of the low quantity of materials available (blood, tissue), or the difficulty to get immune kits for non-model species.

The recent advent of ‘omics’ technologies provides opportunities to explore a wide set of genes and pathways involved in the response to novel conditions experienced at invasion fronts. Genome scans that aim at analysing genome-wide variations in order to detect loci evolving under positive directional selection, have been successfully performed to study some invasion cases (e.g.,^17-19^). They have enabled to detect signatures of contemporary adaptation associated with invasion, some of them being related to immunity ^20^. Nevertheless, this genomic approach based on the detection of single nucleotide polymorphism outliers does not enable to identify the molecular processes driving the rapid adaptation that might occur within a few generations. It also prevents from catching the importance of phenotypic plasticity in invasion success. Transcriptomics, the analysis of gene expression at a genome wide scale, is a complementary approach that might allow filling these gaps and deciphering the molecular mechanisms that underlie phenotypic changes^21,22^. Indeed, gene expression is an essential mechanism for rapid acclimatization or adaptation to novel environments. Transcriptomics has hence been applied to identify genes and gene regulatory pathways that contribute to phenotypic variation at invasion fronts. Evidence for transcriptomic divergence along invasion gradients have already been reported^19,28^, sometimes emphasizing the importance of immune regulation in invasion success ^29^. However there is yet no study combining analyses of immune gene expression and potential variations in parasite pressures along invasion routes. These elements are essential to test the EICA hypotheses and further understand the interactions between immune system regulation and invasion success ^16^.

Here, we examined transcriptional profiles along invasion routes of the house mouse *Mus musculus domesticus* and the black rat *Rattus rattus* currently invading Senegal (West Africa). These two rodent species have been colonizing Senegal eastwards from western coastal colonial cities since the beginning of the 20^th^ century ^30,31^. Our previous study based on few immune effectors highlighted higher inflammatory and/or antibody-mediated responses in recently invaded sites compared with anciently invaded ones for both invaders ^32^. These immune variations could be considered as responses to novel parasite pressures encountered in recently invaded areas. Surveys of bacterial communities in Senegal seemed to corroborate this prediction ^33^ as both quantitative and qualitative changes in bacterial composition were observed along the house mouse and black rat routes of invasion. However, this pattern could not be generalised to any ‘parasite’ pressure. A decrease of the overall prevalence in gastrointestinal helminths was also observed in recently invaded sites compared with anciently invaded ones, suggesting enemy release for these macroparasites ^34^ from introduction areas to invasion fronts. We therefore developed a whole RNA sequencing (*i.e.*, RNAseq) approach to assess without any *a priori* differential gene expression patterns between populations from anciently and recently invaded sites. For both species, we focused on the spleen as it is an immune related tissue in Vertebrates. We tested the null hypothesis that immune gene expression patterns would not differ along invasion routes (i.e. recently invaded sites *vs*. anciently invaded sites). We investigated two alternative hypotheses. On the one hand, we expected an overall higher immune gene expression of rodent populations in recently invaded sites, as a response to novel parasite pressures encountered ^32^. On the other hand, under the EICA-refined hypothesis, we expected lower levels of expression for genes encoding for energetically costly immune pathways (e.g., inflammation) and higher levels of expression for genes encoding for cost-effective immune pathways (e.g., antibody mediated responses), in recently invaded sites. Enemy release of macroparasites and changes in the composition of bacterial communities in recently invaded sites could mediate such trade-offs between energetically costly and cost-effective immune pathways^33,34^.

More specifically, we investigated the following questions: (i) Are there differences in immune gene expression between anciently invaded and recently invaded populations of mice and rats? (ii) Are there functional categories of immune genes that are over-represented among the differentially expressed genes? iii) Do these results support the hypothesis of an overall increase of immune gene expression, or are these results in line with the EICA or EICA-refined hypotheses?

## Results

Results were obtained from the transcriptome analysis of the spleen of two invasive rodent species, the house mouse and the black rat, each one being trapped in four anciently invaded (more than 100 years) and four recently invaded (less than 30 years) sites defined according to historical and longitudinal surveys ^30^.

### Qualitative description of expression patterns

Thirty-two transcriptome libraries were sequenced (see Table 1 for details). They were built from pools of 6 to 10 individual samples for each species and for each site, with two equimolar pools (replicates) per site and per species. These libraries produced an average of 51.6 million read pairs. The numbers of retained reads post-filtering ranged from 37.2 M to 76.7 M, corresponding to 92.82% of the reads. When considering the house mouse data (3 mismatches/read pair allowed), an average of 85.68 % of the quality-filtered reads mapped to the *M. musculus domesticus* reference genome, and 1.58% of these latter mapped to multiple regions. When considering the black rat *R. rattus* data and the possibility for 9 mismatches/read pair with *R. norvegicus* reference genome, 81.66% of the quality-filtered reads were mapped to the reference genome of the related *R. norvegicus*. Among them, 0.98% mapped to multiple regions of the reference genome.

**Table 1:**
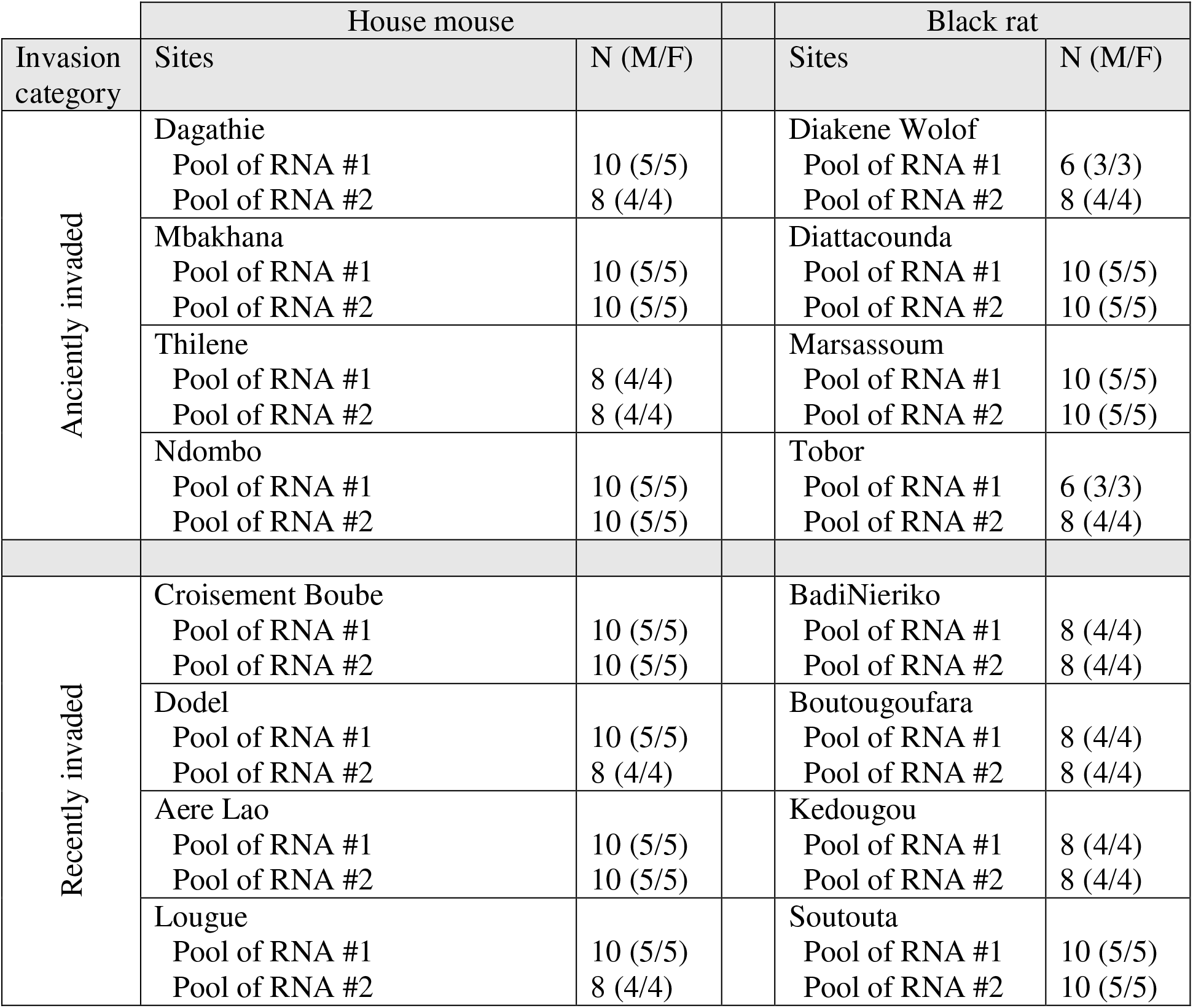
Sampling sites and their invasion-related categorisation as well as rodent sample sizes included in each biological replicate on the house mouse and black rat invasion routes. N: total number of rodents included; M: number of males; F: number of females.

The multidimensional scaling analysis (MDS) performed on the house mouse standardized read counts revealed a slight opposition between sites corresponding to anciently and recently invaded areas on the first axis, except for Aere Lao that did not group with recently invaded sites (Suppl. Fig. S1a). No separation of sites according to invasion-related categories was visible on the MDS performed on the black rat data (Suppl. Fig. S1b).

### Differential expression (DE) analyses

For both datasets, a high dispersion of the total number of reads was observed between biological replicates (Suppl. Fig. S2a,b). Therefore we applied differential expression (DE) analyses using two strategies (see the Methods section). The first one named ‘4vs4 approach’ considered a single randomly chosen biological replicate for each site and analysed the 256 comparisons of the relative log expression (RLE)-transformed normalized read counts between anciently and recently invaded sites. This approach is highly conservative, considering the high variability observed between samples. The second one named ‘8vs8 approach’ considered the eight possibilities of ‘site x replicate’ and employed a generalized linear model (GLM) to take into account the fact that each site was represented by two biological replicates in this analysis. This approach is less conservative but might lead to false positive. In consequence, we only kept the DE genes of the ‘8vs8 approach’ that were also found to be DE in at least 85% of the 256 ‘4vs4 approach’ comparisons.

Along the mouse invasion route, over the 17,853 filtered genes (more than 10 occurrences) tested using edgeR and the ‘4vs4 approach’, 18 genes were always differentially expressed between anciently and recently invaded sites. Over the 16,630 filtered genes (more than 10 occurrences) tested using the ‘8vs8 approach’, we detected 593 differentially expressed genes among which 364 were found in at least 85% of the 256 previous ‘4vs4 approach’ comparisons. Also note that five genes were detected by both ‘4vs4’ and ‘8vs8’ approaches (Hal, Rnf183, Serpina6, Wif1, 9030619P08Rik; see details in Suppl. Table S1a).

Along the black rat invasion route, 54 genes were always differentially expressed between anciently and recently invaded sites among the 13,190 filtered ones (more than 10 occurrences) analysed when using the ‘4vs4 approach’. We found 268 DE genes among the 12,747 filtered ones (more than 10 occurrences) analysed with the ‘8vs8 approach’ among which 83 were detected in at least 85% of the ‘4vs4’ comparisons. Forty-two differentially expressed genes were common to the ‘4vs4’ and ‘8vs8’ approaches. Results are detailed in Suppl. Table S1b.

We have focused further analyses on the 364 and 83 genes that were identified as DE in the ‘8vs8’ approach and detected in at least 85% of the ‘4vs4 ‘comparisons, for the house mouse and black rat, respectively, which we qualified as “robust” genes. The magnitude of expression differences were significantly different between the mouse and rat, with higher ‘log fold changes’ observed for mouse genes (Fig. 1, Wilcoxon test, *p* = 0.01). The log fold change (LogFC) per gene ranged between −9.04 and 2.34 for the mouse and between −5.79 and 4.49 for the rat robust genes. The average coverage varied from −3.16 to 11.94 LogCPM (counts per million) for the mouse and −2.41 to 9.56 for the rat robust genes.

**Figure 1.**
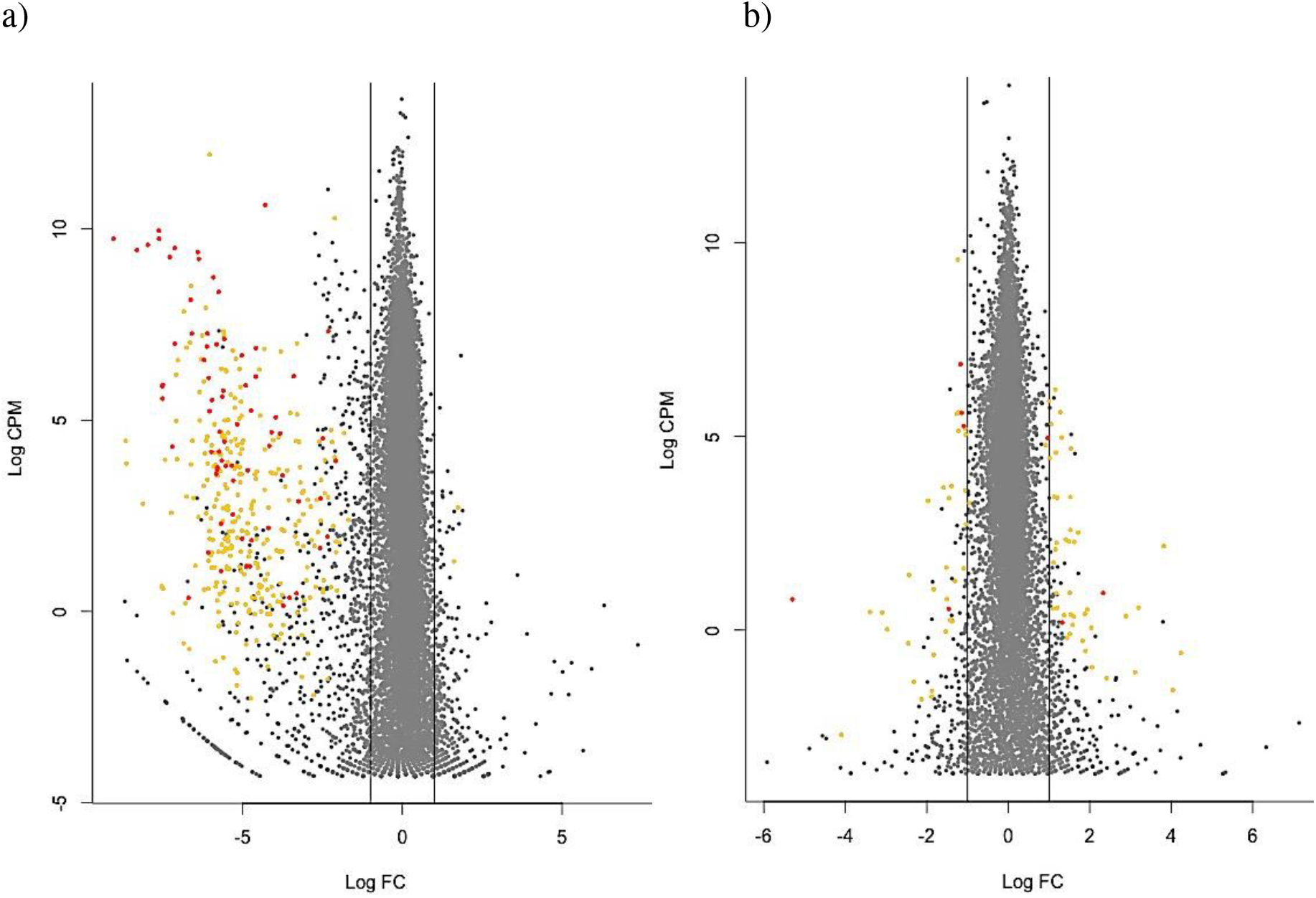
Significantly differentially expressed genes (orange dots) and immune related genes (red dots) between anciently invaded and recently invaded areas along a) the house mouse and b) the black rat invasion roads. The 73 immune related genes belonging to biological processes found to be significantly enriched are represented for the house mouse. Vertical lines indicate 1 log fold change (Log FC). The x-axis indicates genes that are down-(negative values) and up-regulated (positive values) in anciently compared to recently invaded areas. The y-axis represents the average level of gene expression using read counts per million (Log CPM).

### Functional analyses of DE genes

Considering the house mouse dataset, STRING tool provided at least one gene ontology (GO) annotation for 345 of the 364 DE genes (Suppl. Table S1a). Among the 2,098 GO annotations found, 357 GO terms corresponding to biological processes were significantly enriched (FDR < 0.05) when compared to the house mouse reference genome. In particular, 10 immune-related functions were significantly enriched, including inflammatory (GO:0006954), humoral (GO:0002455, GO:0006959), and innate (GO:0045087, GO:0034097) responses or activation/regulation of immunity (GO:0006952, GO:0002253, GO:0050776). We also detected enrichment for two biological processes related to response to wounding (GO:0009611, GO:1903035). These enriched GO terms related to immunity corresponded to 73 DE genes, i.e., 20% of all DE genes (see Suppl. Table S1a). When taking into account redundancies using REVIGO (SimRel semantic similarity measure, similarity cutoff = 0.7), we found 20 main clusters for the category ‘Biological Process’. Inflammation and acute-phase response were part of the second most represented one (Suppl. Fig. S3). Using KEGG algorithm, we detected 47 enriched pathways. We found that 29 immune related genes belonged to these over-represented KEGG biological pathways. The most significant one being the immune ‘Complement and coagulation cascades’ (FDR = 1.56 e-30).

Interestingly, the distribution of average coverage (LogCPM) for these immune-related genes corresponding to enriched GO terms or pathways was shifted towards higher values compared to the 364 robust genes (Wilcoxon test, *p* = 1.52 × 10^−3^; Suppl. Fig. S4). The heatmaps built on the normalized read counts for the immune related genes belonging to the biological processes (as well as pathways) that were found to be over-represented revealed a down-regulation of these genes for all anciently invaded sites and an up-regulation for all recently invaded sites except Aere Lao (Fig. 2, Suppl. Fig. S5a). In this latter population, genes exhibited similar expression patterns than anciently invaded sites.

**Figure 2.**
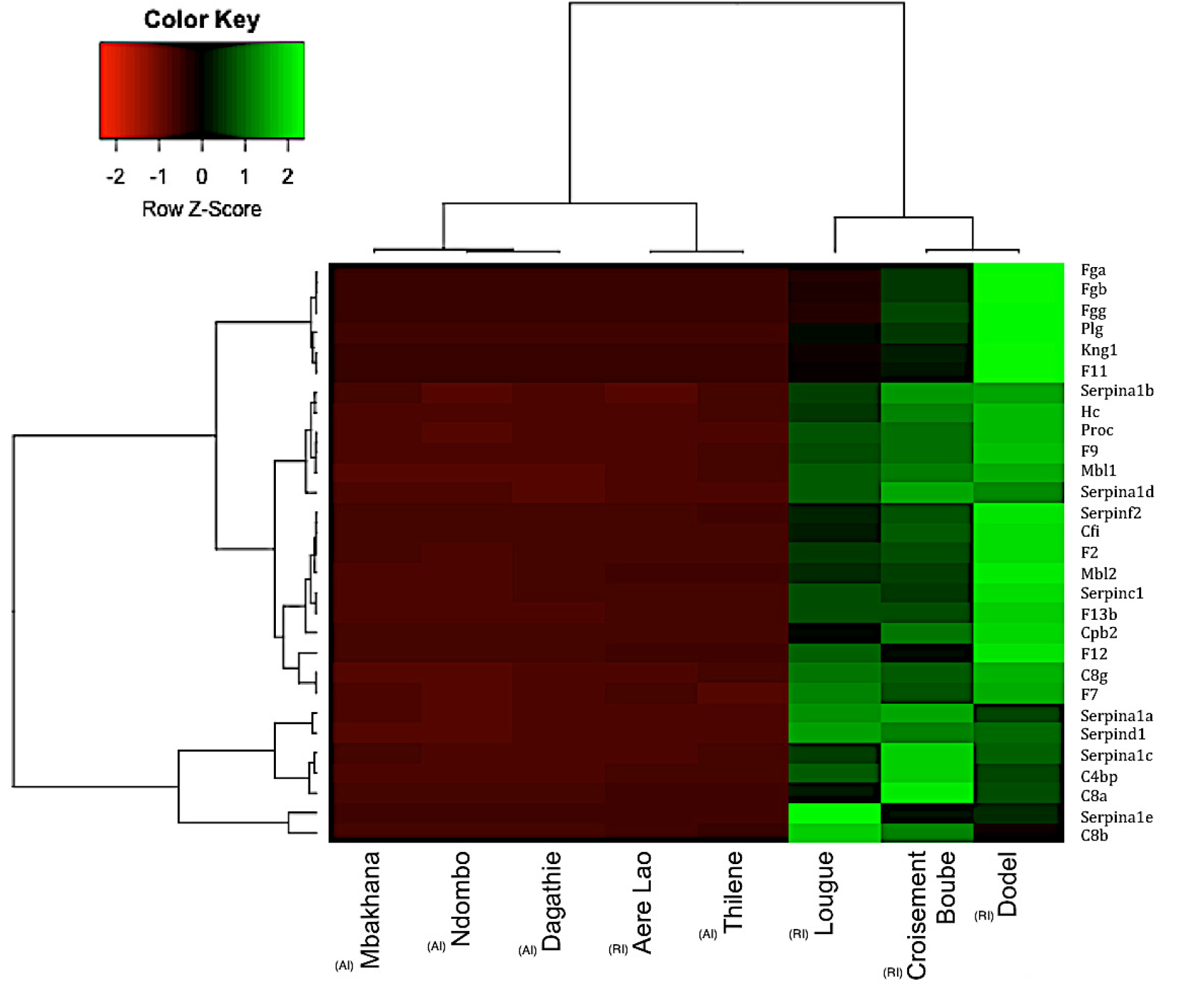
Heatmap of the differentially expressed (DE) genes between the anciently and recently invaded sites of house mouse (*M. musculus domesticus*). The normalized read counts for the expressed genes are shown. For clarity, the heatmap was built in R using heatmap.2 for 29 immune related genes belonging to over-represented KEGG biological pathways. The genes (rows) and samples (columns) were clustered using dendrograms built with Ward distance and hierarchical clustering. Anciently and recently invaded sites are indicated using respectively (AI) and (RI).

Finally, using STRING, we found that the protein-protein interaction (PPI) network based on the 364 DE genes was significantly enriched (*p* < 1.0e-16, 731 edges found, 54 expected), meaning that proteins have more interactions among themselves than what would be expected for a random set of proteins of similar size, drawn from the mouse genome. Two main clusters were detected, one including Cytochrome P450 and UDP glucuronosyltransferase proteins and the other being mainly composed of immune related proteins, including fibrinogen, serin peptidase inhibitor (serpin), apolipoprotein and complement proteins. The PPI interaction network built upon the restricted dataset that included the 73 immune related genes belonging to the biological processes previously found to be over-represented, was also statistically significant (*p* < 1.0e-16, 561 edges found, 13 expected). The network showed a hierarchical structure with the main cluster including the fibrinogen and serine peptidase inhibitor (serpin) proteins, and two other clusters of highly connected proteins including apolipoprotein A and haptoglobin on one hand, and complement proteins on the other hand (Fig. 3).

**Figure 3:**
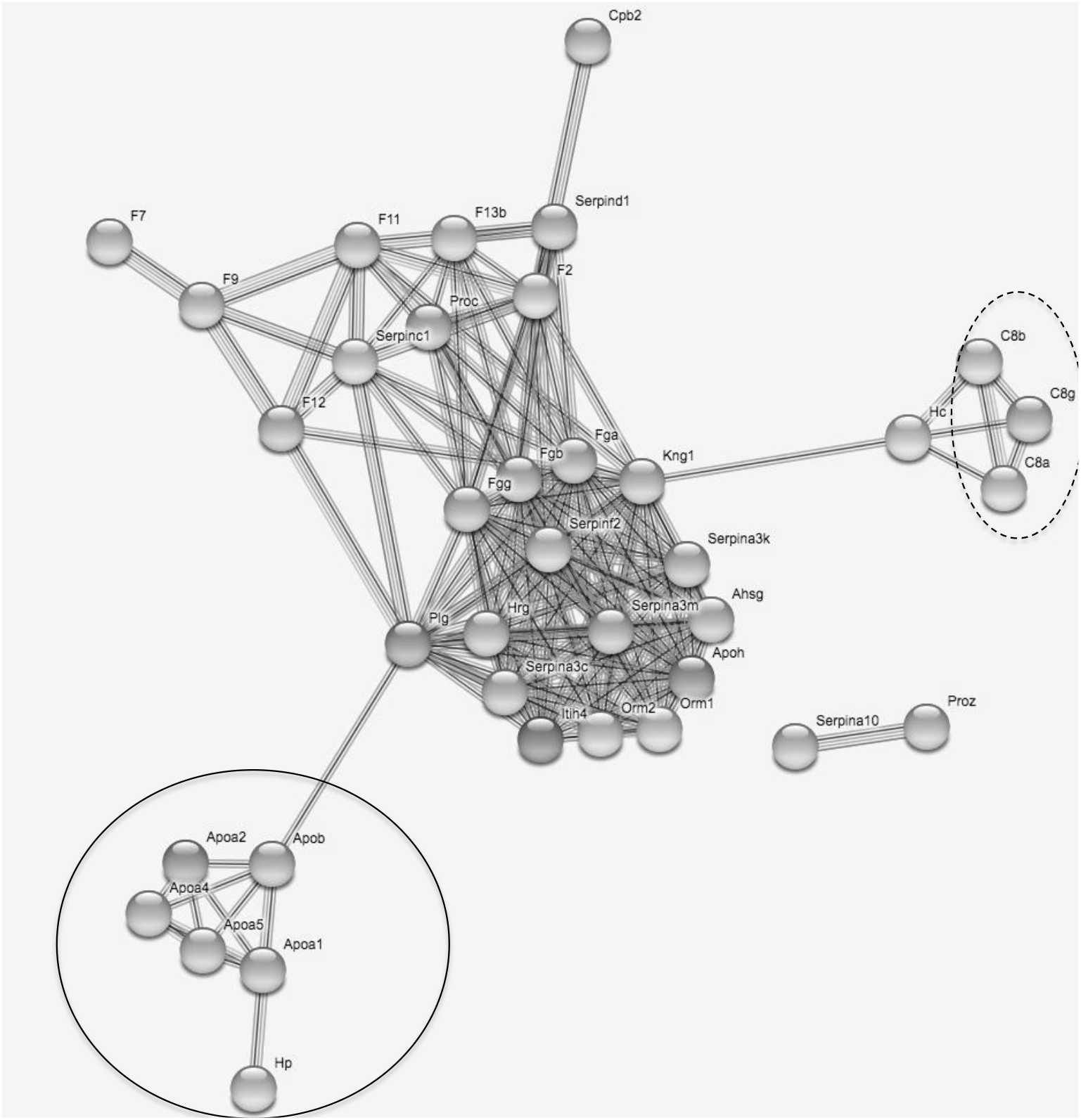
Protein-protein interaction network drawn from the 73 immune genes found to belong to enriched biological processes related to immunity. Nodes correspond to proteins, the thickness of edge network indicates the strength of data support, and the minimum required interaction score was set to 0.7 (high confidence level). The plain circle indicates the alipoprotein A – aloa -and haptoglobin – Hp – protein. The dashed circle indicates the complement proteins (C8). Fibrinogen (Fg) and serine peptidase inhibitor (serpin) proteins are distributed within the highly connected central part of the network.

Considering the *R. rattus* dataset, GO annotations were available for 75 of the 83 DE genes in black rat (Suppl. Table S1b). Eight of these annotated DE genes were related to immune biological processes (i.e., 10.66 % of all DE genes; Suppl. Table S1b). We did not find any biological process or pathway significantly enriched, even when considering redundancies using REVIGO. The protein-protein interaction network built from STRING was not significantly enriched (*p* = 0.09).

## Discussion

In this study, we presented a comparative transcriptomics approach based on spleen RNA sequencing for two invasive rodent species studied along their invasion routes in Senegal. We aimed at identifying adaptive divergence between recently and anciently established rodent populations that may at least partly explain invasion success. More specifically, we investigated differential immune related gene expression that could provide support in favour of the EICA or EICA-refined hypotheses.

Invasion success is supposed to rely on shift in phenotypic traits that would be advantageous at invasion fronts, where species are likely to be exposed to novel environmental conditions ^7,9,35^. Here, using comparative transcriptomics in wild populations, we have evidenced phenotypic shift along the mouse and rat invasion routes in Senegal. Indeed, we detected variations in gene expression levels, a phenotypic characterisation that results from a combination of genotype, environment and genotype–environment interactions ^36^. This phenotypic shift may result from stochastic changes due to population history, including founder events and genetic drift ^7^. Alternatively, it may result from adaptive processes, including phenotypic plasticity^37^ or natural selection^38^. Invasive populations experience reduced genetic diversity due to founder events, but may still evolve rapidly through natural selection from the remaining standing genetic variation or *de novo* mutations ^39^. For black rat and house mouse invasions, the low number of generations that was necessary for both rodents to outcompete native ones after introduction ^30^ suggests however that adaptive variations of gene expression at invasion front might be mediated by phenotypic plasticity ^40,41^

In the house mouse, 20.0% of the genes found to be differentially expressed were annotated with functions related to immunity. We also found that some functional categories of genes were over-represented among these differentially expressed genes. In particular we revealed strong evidence for significantly biased variations in immune gene expression towards an up-regulation of complement and pro-inflammatory cascades in recently invaded sites. These results corroborated our previous functional studies ^32^: immune challenges revealed that antibody-mediated (natural antibodies and complement) and inflammatory (haptoglobin) responses were increased in mice sampled in recently invaded sites compared to those sampled in anciently invaded ones. This study was therefore in line with the hypothesis that parasite-mediated selection could contribute to phenotypic differentiation along invasion routes, potentially in interaction with other environmental factors. However it did not support the EICA or EICA refined hypotheses ^14^. Despite the enemy release patterns detected for mouse helminths in Senegal, from anciently invaded sites to recently invaded ones (decrease of specific richness and of overall prevalence, see ^34^), we did not find any evidence of a global dampening of immune related gene expression in recently invaded sites (the pattern expected according to the EICA hypothesis). We neither observed changes in immune related gene expression that would depend on the relative energetic costs of immune pathways (the pattern expected according to the EICA refined hypothesis). Indeed, inflammation and complement pathways were both up-regulated and they are respectively considered to be energetically costly and cost effective, see ^15^. Therefore the up-regulation of all immune genes found to be differentially expressed in sites recently invaded by the house mouse strongly supported the assumption of an increased overall infection risk in recently invaded sites. Our results previously gathered on the community of pathogenic bacteria in these rodents are in line with this potential increased risk. Indeed, we observed changes in bacterial communities between anciently and recently invaded sites. In particular, we detected introduction of exotic bacterial genera and/or changes in the prevalence of endemic ones, see ^33^. Similar rapid changes in immunity have been observed in other invading organisms, but these studies revealed a wide array of patterns that did not enable to conclude on the potential genericity / specificity of the EICA hypotheses ^16^. We advocate for further studies using wide ranges of immune effectors, including ‘omics methods’ ^42^, to improve our understanding of the relationships between biological invasion and immunity.

Interestingly, the site of Aere Lao did not provide the expected signature along the mouse invasion route, as it provided similar results than anciently invaded sites. The difference of Aere Lao among recently-invaded sites was already noticeable in some of the effectors (complement; haptoglobin) previously measured by immune challenges ^32^. This pattern may not be related to parasitological data, because endemic parasites were found at high levels of prevalence in this site^33,34^. On the contrary, it may reflect a more ancient invasion history of the house mouse in this locality than in the neighbouring ones, for which we have historical data ^30^. Indeed, Aere Lao is a bigger city than Dodel, Croisement Boubé or Lougue (P. Handschumacher, pers. com.) and a market place that may have been colonised earlier by the house mouse.

Concerning *R. rattus*, changes in gene expression levels were of lower amplitude than for the mouse genes and no particular pathway (including immune related ones) was significantly over-represented. This absence of clear pattern suggested that stochastic events may have strongly shaped the phenotypic variations observed along *R. rattus* invasion route. However, this conclusion must be taken cautiously as bioinformatics limitations could underlie this result (transcriptome annotation was less efficient for *R. rattus* than for *M. musculus domesticus* as we used *R. norvegicus* transcriptome as a reference). The eight immune related genes found to be differentially expressed encode for proteins that interact with bacteria (*Rnase6*, *Ptprz1*), viruses (*Ltc4s*, *Hspa1b*) or parasites (*Gfi1b*). Some of them were down-regulated in recently invaded sites (e.g., *Rnase6)* while others were strongly up-regulated (e.g., *Ptprz1*). So, no specific pattern emerged with regard to potential variations in infection risks. Therefore variations of immune phenotypes, as reflected by gene expression in spleen, were unlikely to be the main driver of the black rat invasion success (considering that invasion success relies on the rapid shift of phenotypic traits in newly colonized areas and not on pre-adaptation in the anciently invaded range). Interestingly, we also previously found from a functional immune study that differences in immune responses between the rat anciently and recently invaded sites were less marked than for the mouse ^32^. In addition, we could hardly identify other biological processes that could reflect life history traits adaptation with regard to invasion. Those processes that were found to be associated with differentially expressed genes covered a wide range of functions including clustering of voltage sodium channels, negative regulation of cell-matrix adhesion and protein dephosphorylation and response to cyclic Adenosine Monophosphate (cAMP). This latter function could be interesting with regard to the rat invasion success as cAMP is central to the regulation of insulin and glucagon secretion ^43^. Variations in cAMP gene expression could therefore mediate differences in response to stressful situations, including starvation or fight-or-flight response ^44^. It would be interesting to analyse whether such differences could result in different behavioural phenotypes between rats trapped in anciently invaded sites (expected to exhibit little performance and high stress response in novel environments) and recently invaded ones (expected to exhibit high response capacity in novel environments) ^45^. Therefore it seems even more important to further perform transcriptomics analyses on other organs and tissues to identify the phenotypic changes and ecoevolutionary processes linked with the black rat invasion success. Brain, where key genes underlying behavioural invasion syndrome are expected to be expressed, could be a relevant organ to target in the future ^46^.

In conclusion, our work revealed changes in transcriptomic profiles along invasion routes for both the house mouse and the black rat. It is likely that different processes, including colonization history and/or alternative mechanisms by which species adapt to novel environments, may have mediated the invasion success of these two rodent species. The patterns observed could potentially be driven by increased infection risks in recently invaded sites for the house mouse and by stochastic changes for the black rat. It could be interesting to take into account the potential variability of these patterns with regard to the targeted tissue or organ. We here focused on rodent spleen. Results would perhaps have been different if we had targeted other lymphoid organs, lymphatic tissues or non-immune targets. In the future, genomic studies and experimental work^19,22^ could enable to decipher whether differences in gene expression were driven by phenotypic plasticity or directional selection during or after invasion, or reflected the colonization history of rodents. Moreover, it would be important to validate whether these changes in phenotypic traits influenced ecological dynamics e.g., ^47^ and in turn, invasion success.

## Methods

### Ethical statements

Sampling campaigns within private properties were realized with the prior explicit agreement from relevant familial, local and institutional authorities. None of the rodent species investigated here has protected status (see list of the International Union for Conservation of Nature). All animal-related procedures were performed according to official ethical guidelines provided by the American Society of Mammalogists ^48^. All protocols presented here were realized with prior explicit agreement from relevant institutional committee (CBGP: D 34-169-1). They were carried out in accordance with requirements of Senegalese and French legislations.

### Sampling sites, sample collection and RNA extraction

We used some of the spleen samples of *M. m. domesticus* and *R. rattus* that were previously analysed for the bacterial metabarcoding study previously described ^33^. Briefly, for both invasive species, rodent trapping was performed in four anciently invaded (more than 100 years) and four recently invaded (less than 30 years) sites defined according to historical and longitudinal surveys ^30^. Autopsies were realized at approximately the same period of the day for all individuals (11 am – 14 pm). Spleens were collected and immediately placed in RNAlater, stored at 4 °C overnight then at −20 °C until further analyses.

Twenty adult rodents per site were considered excepted for five sites where only 14-18 samples were available (see Table 1). Total RNA was extracted from approximately 5mg of spleen for each sample using AllPrep 96 DNA/RNA Kit (Qiagen). The quality and quantity of the purified RNA was assessed by gel electrophoresis and NanoDrop spectrophotometer (Thermo Scientific) before pooling, and Bioanalyzer 2100 (Agilent) after pooling (see below). For each rodent species, sixteen equimolar pools (2 per sites) that combined 6 to 10 individual samples with equilibrated sex ratio were made.

### cDNA library preparation and RNA sequencing

cDNA library construction, sequencing and sequence alignment of the filtered reads were performed at MGX platform. RNA-Seq libraries were constructed with the Truseq stranded mRNA sample preparation (Low throughput protocol) kit from Illumina. One microgram of total RNA was used for the construction of the libraries. The first step in the workflow involved purifying the poly-A containing mRNA molecules using poly-T oligo attached magnetic beads (Poly-A based mRNA enrichment). Following purification, the mRNA was fragmented into small pieces using divalent cations under elevated temperature. The cleaved RNA fragments were copied into first strand cDNA using Actinomycine D, random hexamer primers and SuperScript II reverse transcriptase for *M. m. domesticus* samples or SuperScript IV reverse transcriptase for *R. rattus* samples. The second strand cDNA was synthesized by replacing dTTP with dUTP. These cDNA fragments then have the addition of a single ‘A’ base and subsequent ligation of the adapter. The products were then purified and enriched with 15 cycles of PCR. The final cDNA libraries were validated and quantified with a KAPA qPCR kit.

For *M. m. domesticus* samples, four DNA libraries per sequencing line were pooled in equal proportions, denatured with NaOH and diluted to 7 pM before clustering. Cluster formation on a flowcell V3, primer hybridisation and single-end read 50 cycles sequencing were performed on cBot and HiSeq2000 (Illumina, San Diego, CA) respectively.

For *R. rattus* samples, four DNA libraries per sequencing line were pooled in equal proportions, denatured with NaOH and diluted to 8 pM before clustering. Cluster formation on a flowcell V4, primer hybridisation and single-end read 100 cycles sequencing were performed on cBot and HiSeq2500 (Illumina, San Diego, CA) respectively. We performed single-end read 100 cycles for *R. rattus* as up to known the only reference genome available for rats is from another species *Rattus norvegicus*.

### Transcriptome analysis

#### Sequence alignment and gene quantification

Image analyses and base calling were performed using the Illumina HiSeq Control Software and Real-Time Analysis component. Demultiplexing and production of Fastq files were performed using Illumina’s conversion software (CASAVA 1.8.2 for the house mouse data, BCL2FASTQ 2.17 for the black rat data). The quality of the raw data was assessed using FASTQC from the Babraham Institute and the Illumina software SAV (Sequencing Analysis Viewer).

A splice junction mapper, TOPHAT ^49^ (v2.0.9 for the mouse data, v2.0.13 for the black rat data), using BOWTIE ^50^ (v2.1.0 for the mouse data, v2.2.3 for the black rat data), was used to align the reads to the *M. musculus* genome (UCSC mm10) or the *Rattus norvegicus* genome (UCSC rn4) with a set of gene model annotations (genes.gtf downloaded from UCSC on March 6, 2013 for the mouse genome, on July 17, 2015 for the brown rat genome). Final read alignments having more than 3 mismatches (house mouse) or 9 mismatches (black rat) were discarded. Again we used a different threshold for mouse and rat data as the reference genome available for this latter is from *Rattus norvegicus*. Read alignment rates were above 83,3 % for all *M. m. domesticus* libraries and 74.0 % for all *R. rattus* libraries.

Samtools (v0.1.18 for the mouse data, v1.2 for the black rat data) was used to sort the alignment files. Then, gene counting was performed with HTSEQ count (v0.5.4p5 for the mouse data, v0.6.1p1 for the black rat data) using the union mode ^51^. The data is from a strand-specific assay, the read has to be mapped to the opposite strand of the gene.

#### Analysis of differential gene expression between populations

Differentially expressed (DE) genes were identified using the BIOCONDUCTOR ^52^ package edgeR v3.4.0, under R 3.0.2 for the mouse data, v3.8.6 under R 3.2.3 for the black rat data, ^53^. Data were normalized using the relative log expression Rle normalization factors ^54^. The design of the experience includes two statistical factors: the main factor is the date of the invasion (two levels: recently and anciently invaded), and the second factor is the locality in which the rodent has been sampled (eight localities in total: four for the recently invaded and four for the anciently invaded). In each locality, two replicate samples were prepared (for example, there are two samples for the locality Dagathie). In order to take into account this complex experimental design, two analytical approaches have been used to identify the differentially expressed genes. First, the decision has been taken to keep only one out of the two replicates per locality. This allows to suppress the second factor (locality) of the analysis: we compare the four recently invaded versus the four anciently invaded regions, and here the four samples per condition are independent from each other and considered as replicate (‘4vs4 approach’). As there is no reason to keep the first or the second locality replicate, all the 256 combinations of replicate selection has been made. As an example, the first combination is: (Da1, Mb3, Th5, Nd7) vs (Cr9, Do11, Ae13, Lo15). Secondly, the eight samples from each condition were kept, and the locality factor was added to the design in the statistical analysis settings in order to consider that replicates were paired by locality (‘8vs8 approach’). Before statistical analysis, genes with less than 10 occurrences, cumulating all the 8 or 16 analysed samples per species, were filtered and thus removed. It enabled to limit the number of statistical tests and therefore to lower the impact of multiple tests corrections. In these two approaches, the *p*-value threshold was set to 5%, after application of the Benjamini-Hochberg method for multiple testing correction.

As a high variability is observed between the samples, combining the results of these two approaches seemed to be a prudent choice, enabling the results to be as robust as possible. Thus, the genes identified with the ‘8vs8 approach’ and declared as DE in more than 85% of the 256 combinations of the ‘4vs4 approach’ were eventually declared as DE. The threshold of 85% were set according to the visualisation of a barplot representing the number of ‘4vs4’ comparisons in which a gene (highlighted in the ‘8vs8 approach’) is found to be differentially expressed.

We performed a multidimensional scaling analysis (MDS based on log fold change distance) on normalized read counts of expressed genes (excluding weakly expressed ones, see above) using R v3.0.1 ^55^ and the edgeR libray (plotMDS function). It enabled to visualize distances in overall gene expression between sites.

### Functional and enrichment analyses

*M. m. domesticus* and *R. rattus* genes were functionally annotated with gene ontology (GO) terms using Search Tool for the Retrieval of Interacting Genes/Proteins (STRING) v10.5 tool^56,57^. We performed gene enrichment analyses using Fisher’s exact tests on the house mouse and black rat declared DE gene sets using house mouse *M. musculus* and brown rat *R. norvegicus* GO as the background and the software available in STRING. *p*-values were corrected for multiple testing using the Benjamini and Hochberg false discovery rate (FDR). GO terms with a FDR *p*-value < 0.05 were considered as statistically significantly different. The redundancies of significantly enriched GO terms were reduced using REVIGO ^58^ with a similarity cutoff of 0.7. Similarly we performed enrichment analyses on biological processes and biological pathways supplied by KEGG (http://www.genome.jp/kegg/pathway.html), implemented in STRING.

Finally we performed a global protein network analysis based on DE genes using STRING database that included interaction databases (including direct (physical) as well as indirect (functional) associations), genetic interactions and shared pathway interactions. It enabled to show how key components of different pathways interact. Two PPI networks were constructed by mapping all DE genes and immune related DE genes, respectively, to STRING database with confidence score >0.7 (high level). PPI networks were visualized and analysed in CYTOSCAPE software through the web interface of STRING. Enrichment in these protein-protein interactions were tested using Fisher’s exact tests and correction for multiple testing (FDR).

## Data accessibility

All sequence data, including raw reads and assemblies have been deposited on GEO. Gene ontologies for DE genes are provided in Supplementary Tables 1 and 2. Differential expression data are available from the corresponding author on request.

## Acknowledgements

We thank Khalilou Bâ, Mamoudou Diallo, Mamadou Kane, Laurent Granjon, Youssoupha Niang, Aliou Sow and Philippe Gauthier for their help in collecting samples, and J.-M. Duplantier for invaluable advice regarding field work and rodent invasion history. We are indebted to all the Senegalese people who allowed us to trap rodents in their homes. This work was supported by the ANR ENEMI project (ANR-11-JSV7-0006), and by the French Embassy in Senegal, which provided the funding for PhD scholarships. This preprint has been reviewed and recommended by Peer Community In Ecology (https://dx.doi.org/10.24072/pci.ecology.100011).

## Supplementary material

**Figure S1.**
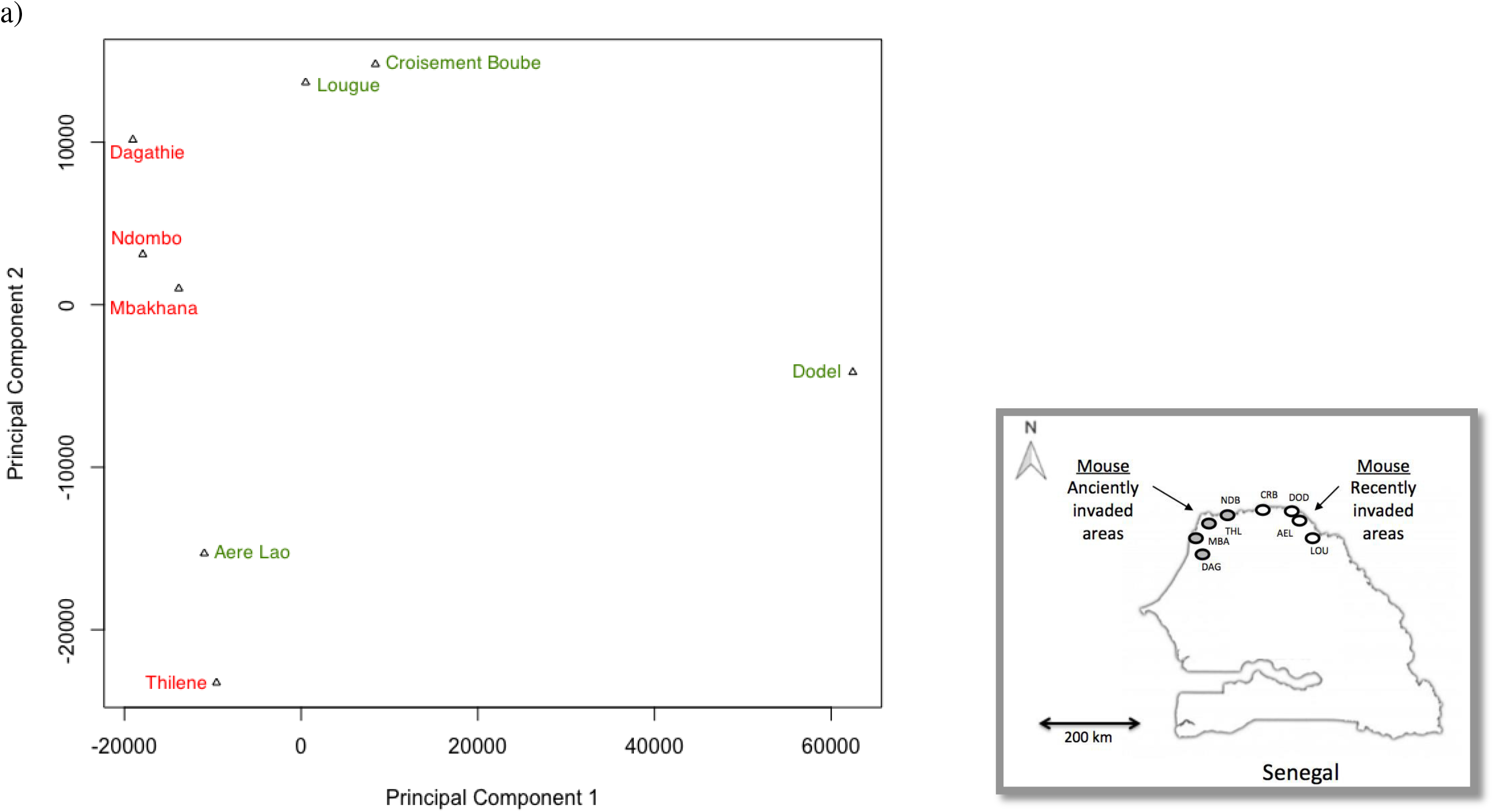
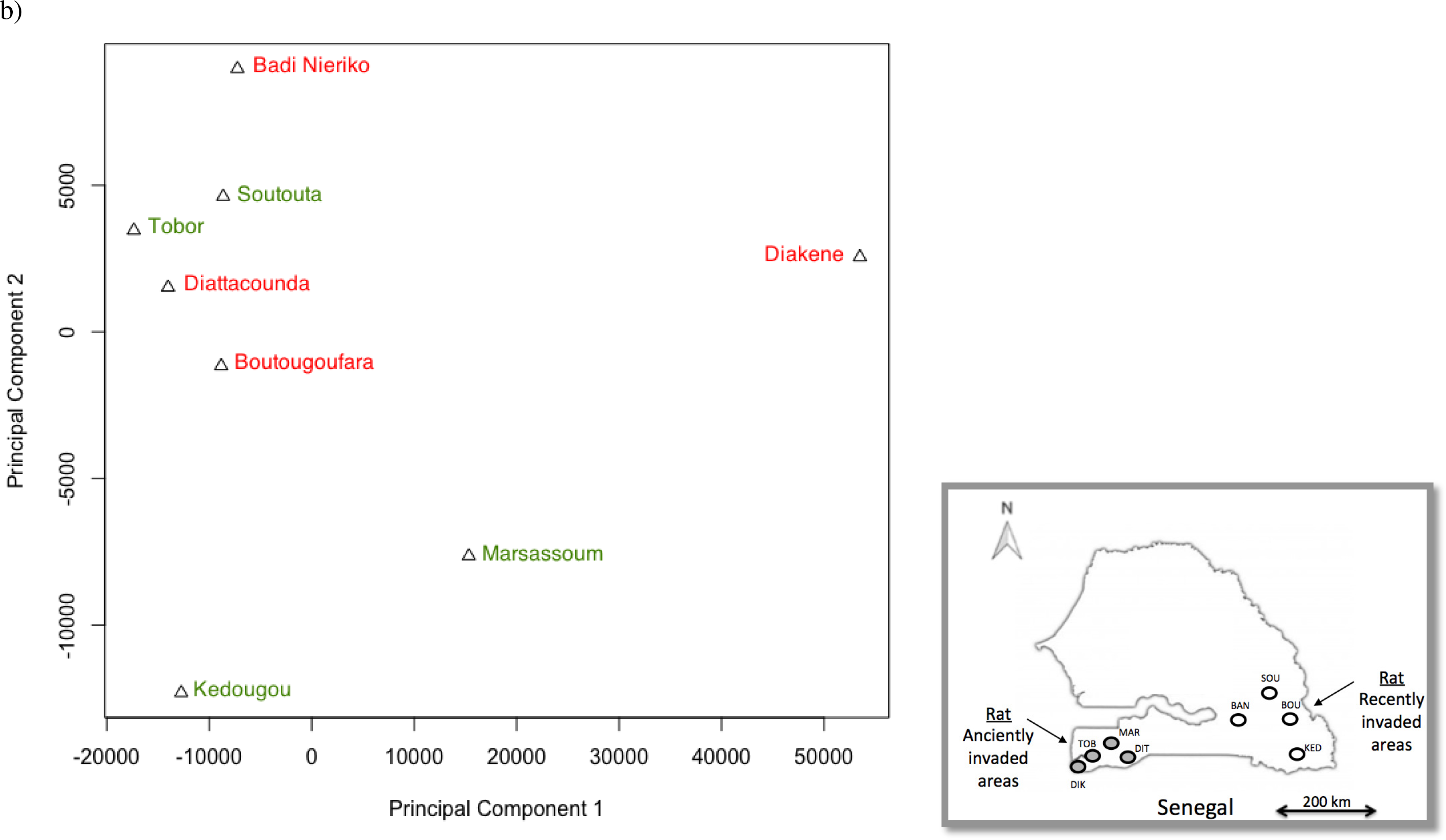
Multidimensional scale (MDS) analyses of RLE-normalized read counts among samples partitioned by sites for a) *Mus musculus domesticus* (16,630 genes) and b) *Rattus rattus* (12,747 genes) invasion routes in Senegal. A map describing sampling sites in Senegal is provided for *M. m. domesticus* and *R. rattus*. Da=Dagathie; Mb=Mbakhana, Th=Thilene, Nd=Ndombo, Cr=Croisement Boube, Do=Dodel, Ae=Aere Lao, Lo=Lougue. Bad=Badi Nieriko, Bou=Boutougoufara, Diak=Diakene Wolof, Diat=Diattacounda, Ked=Kedougou, Mar=Marsassoum, Sou=Soutouta, Tob=Tobor.

**Figure S2.**
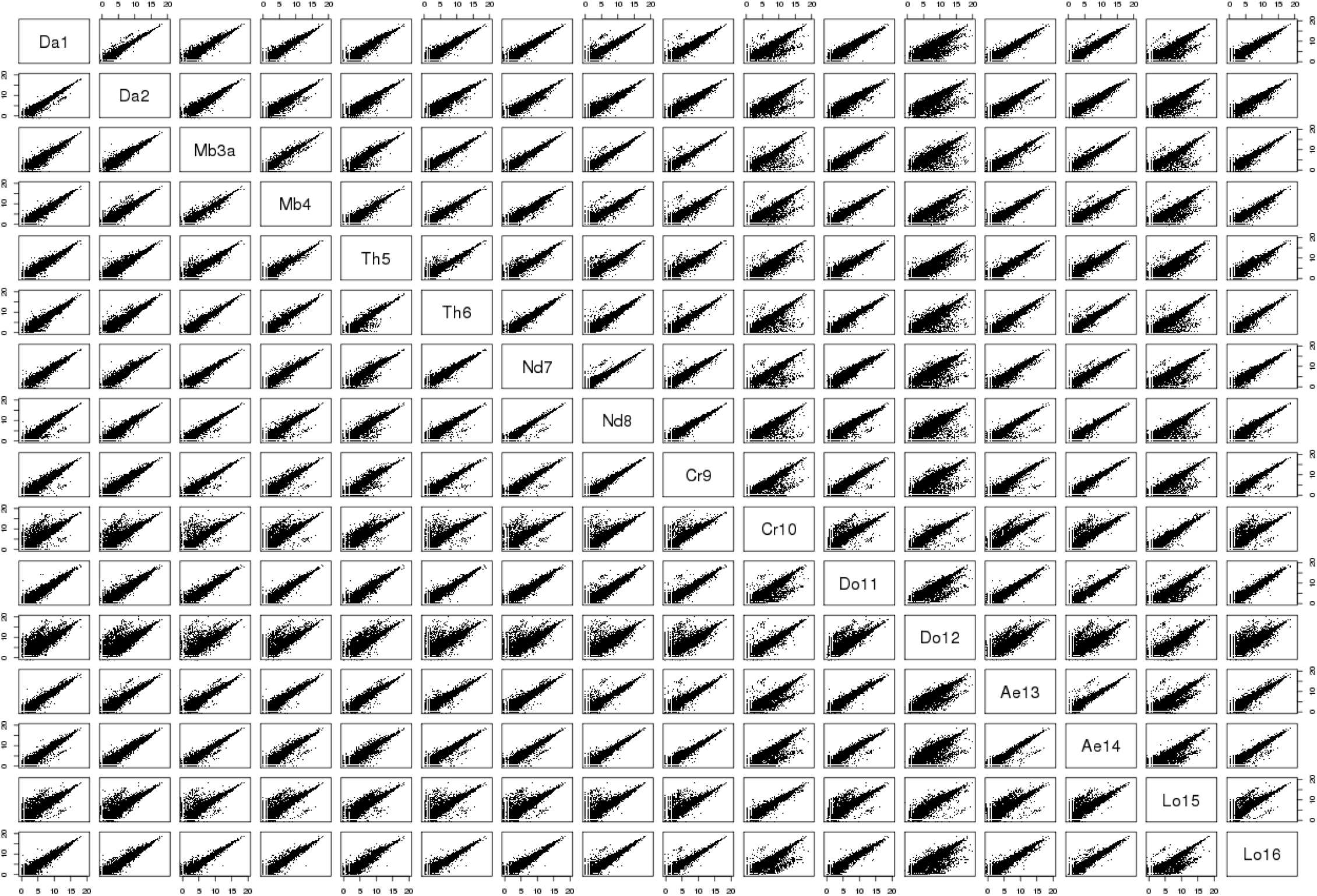
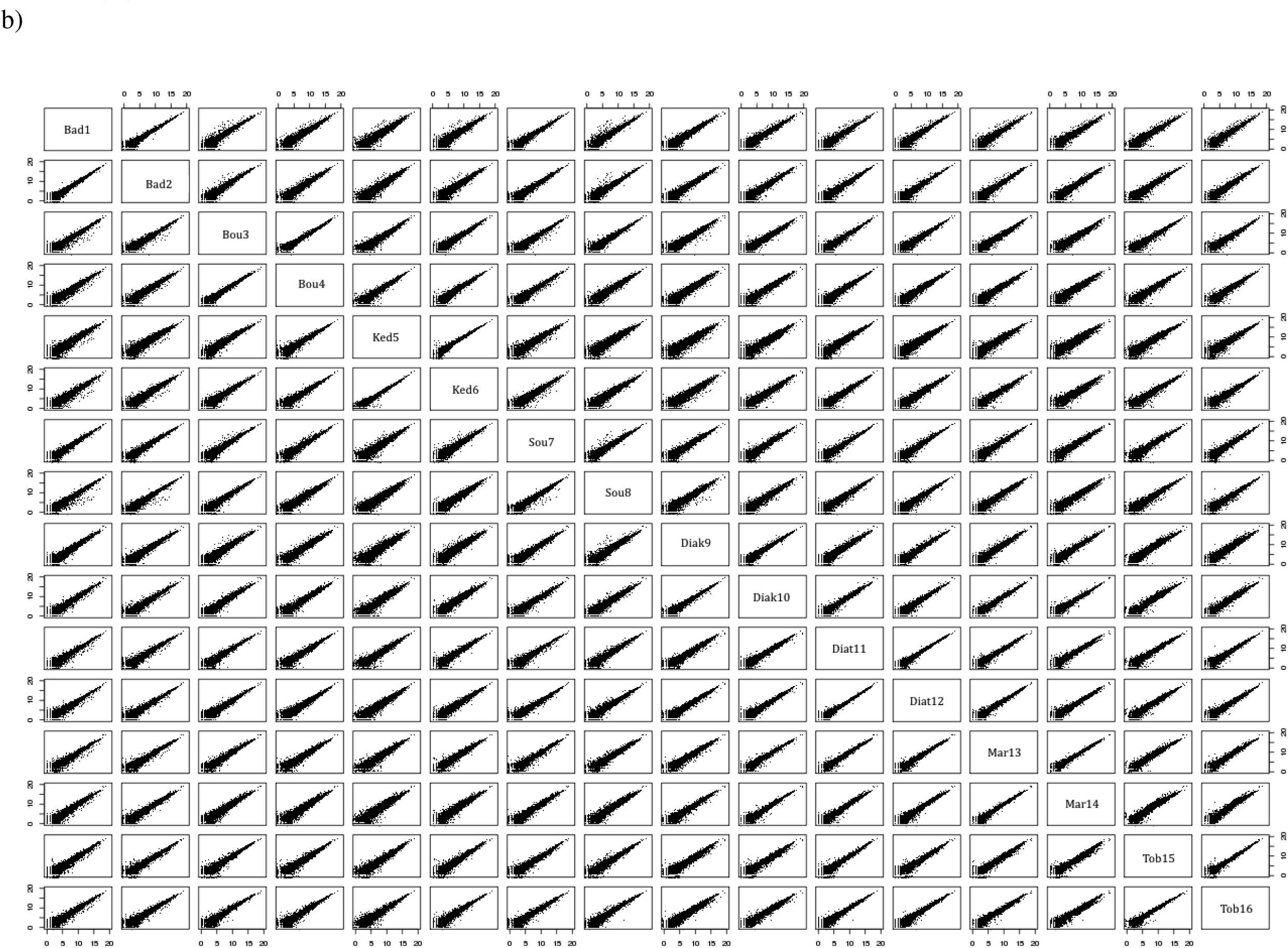
Pairwise comparisons of the RLE-normalized total number of reads between libraries for a) the house mouse and b) the black rat. For the house mouse, anciently invaded sites are: Da=Dagathie; Mb=Mbakhana, Th=Thilene, Nd=Ndombo; recently invaded sites are Cr=Croisement Boube, Do=Dodel, Ae=Aere Lao, Lo=Lougue. For the black rat, anciently invaded sites are: Diak=Diakene Wolof, Diat=Diattacounda, Mar=Marsassoum, Tob=Tobor; recently invaded sites are: Bad=Badi Nieriko, Bou=Boutougoufara, Ked=Kedougou, Sou=Soutouta.

**Figure S3.**
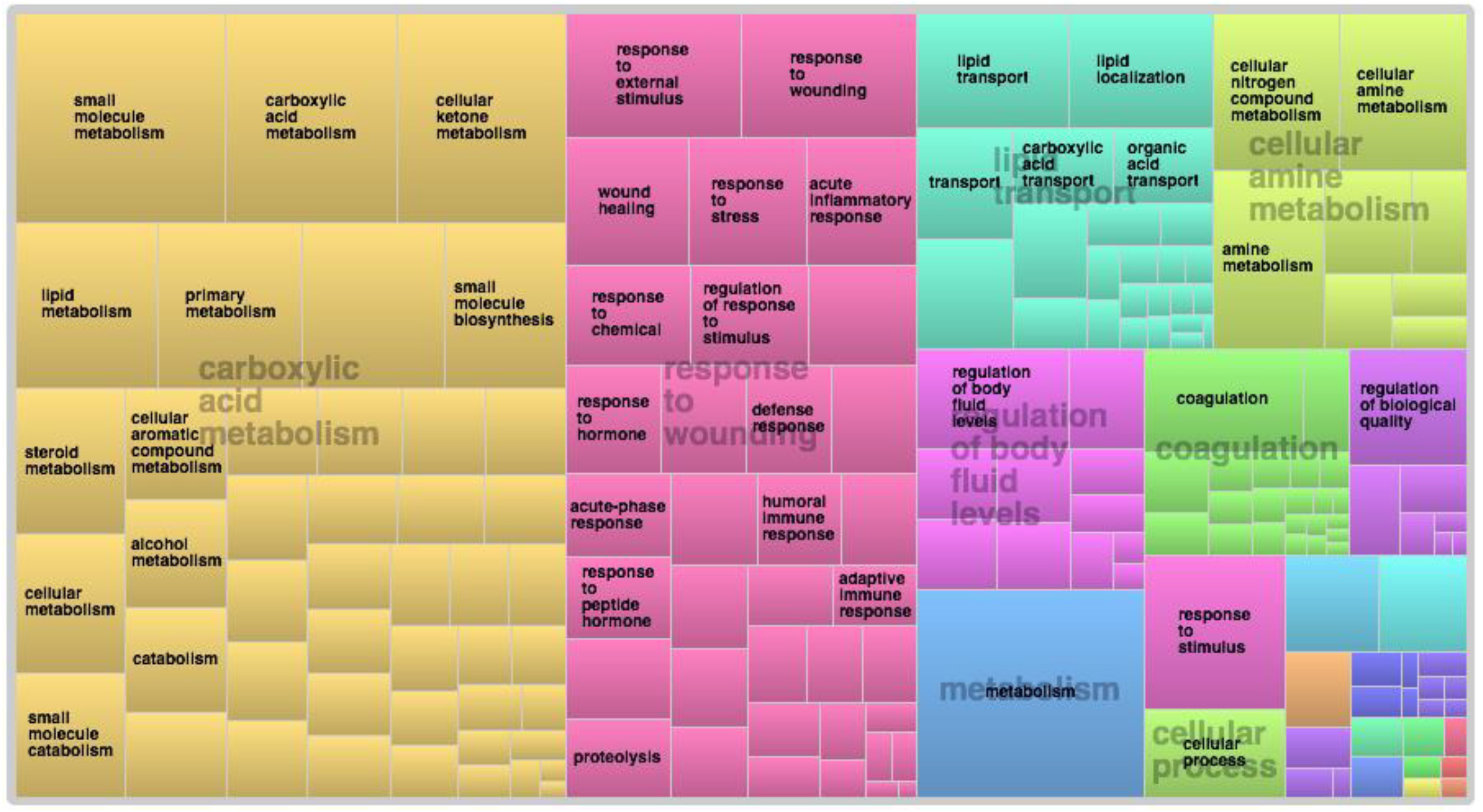
TreeMap view of REVIGO Biological process analyses for the house mouse differentially expressed (DE) genes. Each rectangle represents a single cluster, that are grouped into ‘superclusters’ of related terms, represented with different colours. The size of the rectangles reflects the frequency of the Gene Ontology (GO) term in this set of DE genes.

**Figure S4.**
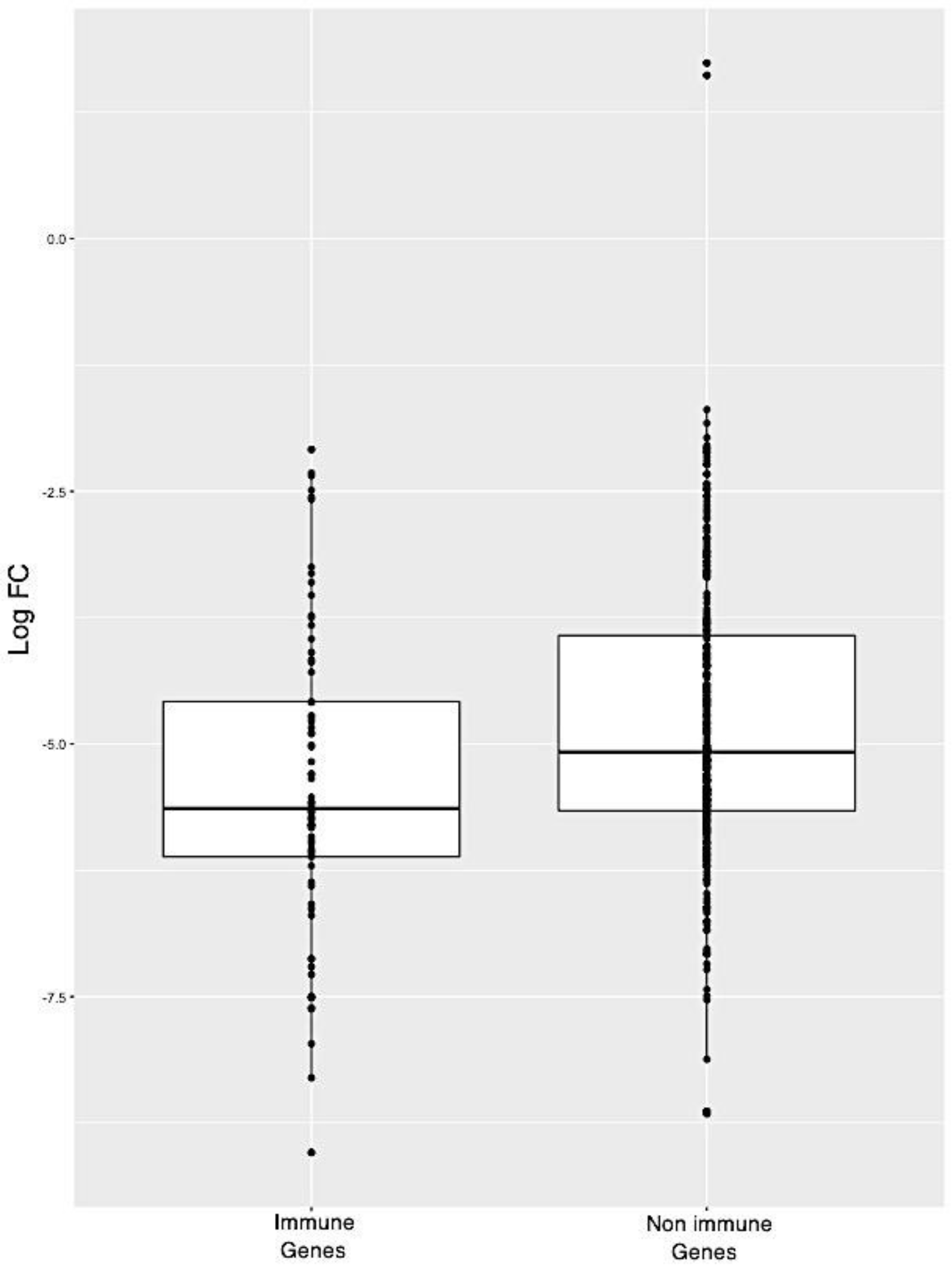
Boxplot representing the level of differential expression (in log fold change (log FC)) for immune related and non immune related robust genes identified along the mouse invasion road.

**Figure S5.**
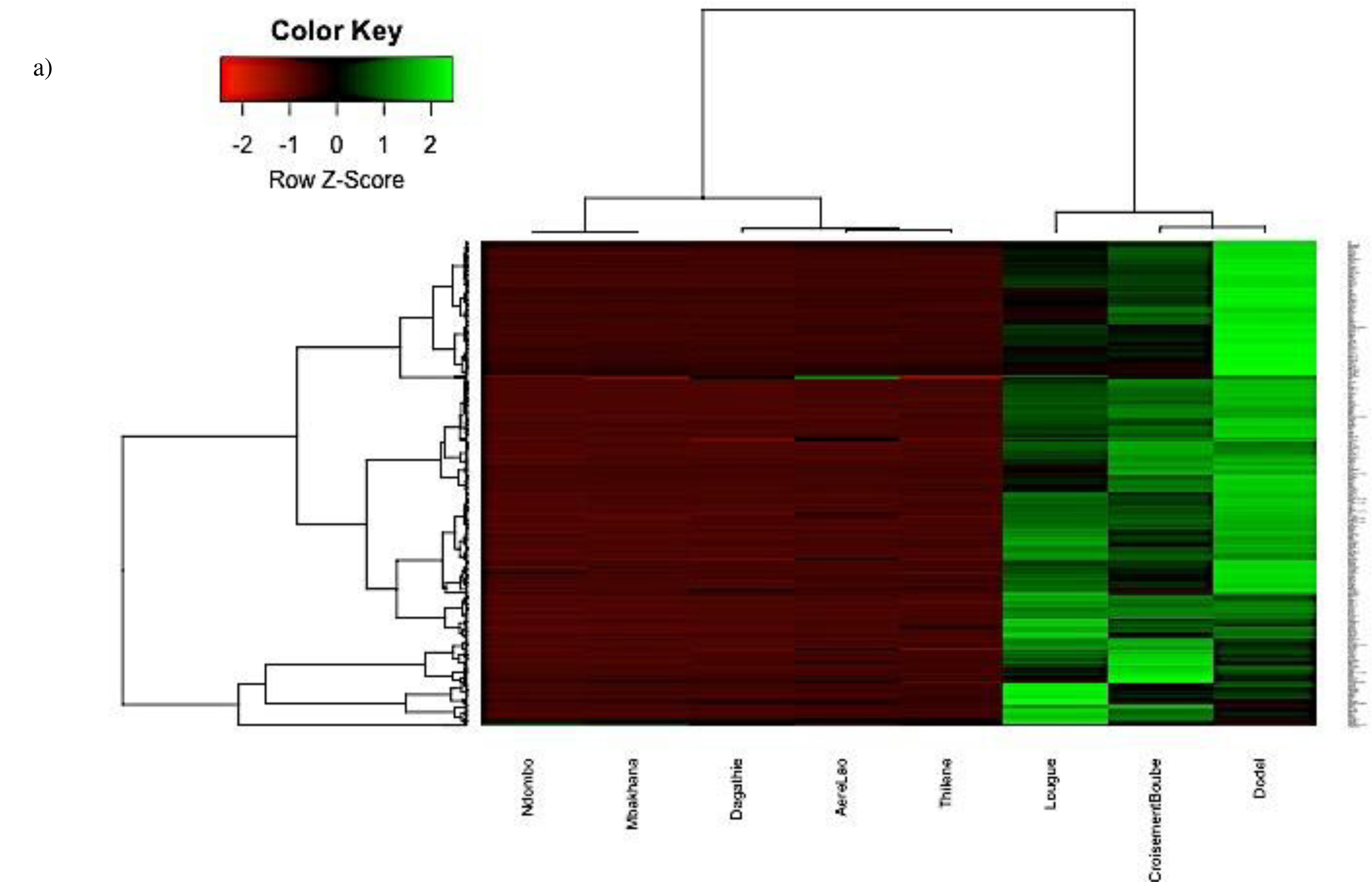
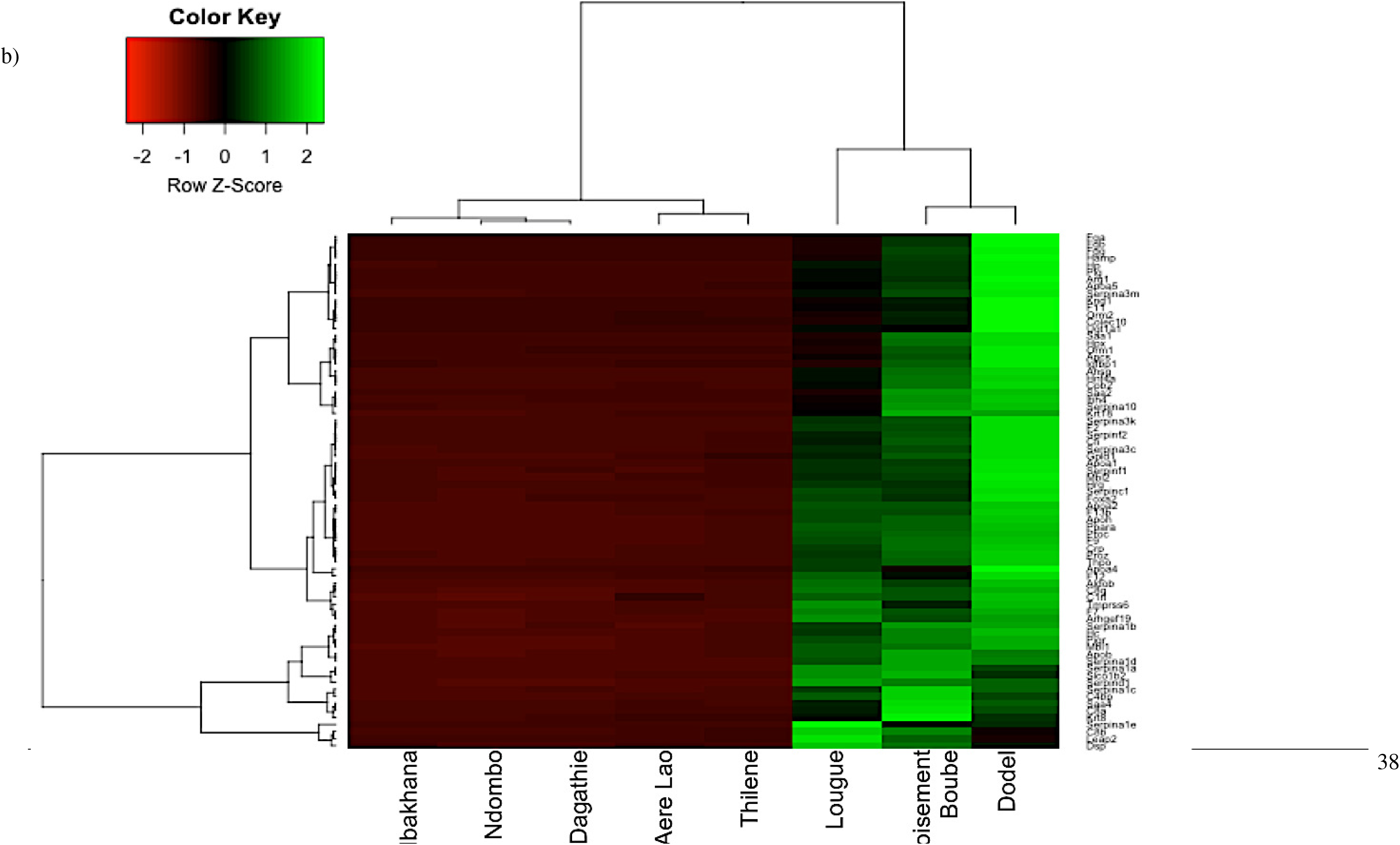
Heatmap of the differentially expressed (DE) genes between the anciently and recently invaded sites of house mouse (*M. musculus domesticus*). The normalized read counts for the expressed genes are shown. Heatmap was built in R using heatmap.2 for a) all 364 DE genes and b) 73 immune related genes belonging to over-represented biological processes. The genes (rows) and samples (columns) were clustered using dendrograms built with Ward distance and hierarchical clustering.

**Table S1.** a) Details of the 18 genes found to be differentially expressed between the anciently and recently invaded sites of *M. musculus domesticus* invasion route using the 4vs4 approach. Genes indicated in bold were found to be differentially expressed with both 8vs8 and 4×4 approaches. b) Details of the 364 genes found to be differentially expressed between the anciently and recently invaded sites of *M. musculus domesticus* invasion route using the 8vs8 approach and found in 85% of the comparisons made using the 4×4 approach. The 73 genes indicated in red are related with immunity and have over-represented GO annotations. The 29 genes underlined and in italics correspond to the immune-related genes with over represented Kegg biological pathways. GO ID corresponds to Gene ontology annotations.

**Table S2.** a) Details of the 54 genes found to be differentially expressed between the anciently and recently invaded sites of *R. rattus* invasion route using the 4vs4 approach. Genes indicated in bold were found to be differentially expressed with both 8vs8 and 4vs4 approaches. b) Details of the 83 genes found to be differentially expressed between the anciently and recently invaded sites of *R. rattus* invasion route using the 8vs8 approach and found in 85% of the comparisons made using the 4vs4 approach. Genes indicated in bold were found to be differentially expressed with both 8vs8 and 4vs4 approaches. Genes indicated in red are related with immunity. GO ID corresponds to Gene ontology annotations.

